# SMURF: embedding single-cell RNA-seq data with matrix factorization preserving self-consistency

**DOI:** 10.1101/2022.04.22.489140

**Authors:** Bingchen Wang, Juhua Pu, Lingxi Chen, Shuai Cheng Li

**Author notes:** These authors contributed equally to this work (Bingchen Wang, Juhua Pu).

## Abstract

The advance of single-cell RNA-sequencing (scRNA-seq) sheds light on cell-specific transcriptomic studies of cell developments, complex diseases, and cancers. Nevertheless, scRNA-seq techniques suffer from “dropout” events, and imputation tools are proposed to address the sparsity. Here, rather than imputation, we propose a tool, SMURF, to embed cells and genes into their latent space vectors utilizing matrix factorization with a mixture of Poisson-Gamma divergent as objective while preserving self-consistency. As for wet lab dataset evaluation, SMURF exhibited feasible cell subpopulation discovery efficacy with the latent vectors on all the eight-cell line mixtures. Furthermore, SMURF can embed the cell latent vectors into a 1D-oval and recover the time course of the cell cycle. SMURF can also serve as an imputation tool, the *in silico* data assessment shows that SMURF paraded the most robust gene expression recovery power with low root mean square error and high Pearson correlation. Moreover, SMURF recovers the gene distribution for the WM989 Drop-seq data. SMURF is available at https://github.com/deepomicslab/SMURF.

## Introduction

The emergence of high throughput single-cell RNA-sequencing (scRNA-seq) flourishes the cell-specific transcriptomes applications (1–3), including cell cycles (4–6), complex diseases (7–9), and cancers (6, 10–14).

scRNA-seq allows us to obtain a gene-cell count matrix, where the rows represent the genes and the columns signify the cells. Nevertheless, this technique has a drawback - it could only catch a small portion of genes during the sequencing process, and such missing entries will be recorded as zero into the count matrix. We call this phenomenon the “dropout” event and such matrix the zero-inflated matrix (15). As such, a zero value in a zero-inflated count matrix contains two possibilities - the first is that the gene expression in a cell is in fact zero, and the second is that it is a fake zero caused by “dropout” events. The sparse gene-cell expression profile might influence the accuracy of downstream analyses and mask potential biological insights (16). Thus, we need to determine the two possibilities concerning zero entries and amend the missing values from false zero to a more statistically realistic value. In computational biology, we termed this data wrangling process as “imputation”.

With the intent of making it more appropriate and valuable for a variety of downstream purposes such as clustering, cell type recognition, dimension reduction, differential gene expression analysis, identification of cell-specific genes, and reconstruction of differentiation trajectory on zero-inflated single-cell gene expression data (16), several scRNA-Seq imputation tools have developed. A systematic benchmarking of 18 scRNA-Seq imputation methods suggested that MAGIC (17), kNN-smoothing (18), and SAVER (19) were state-of-the-art tools (20). MAGIC conducted imputation leveraging information across similar cells with data diffusion (17). Namely, kNN-smoothing imputes the missing entries by collapsing gene-specific UMI counts from the k nearest neighbors (18). SAVER estimates the gene expression level by borrowing information across similar genes utilizing the Bayesian approach (19). Also, in previous work (21), we introduced a self-consistent method I-Impute and demonstrated its robustness compared to SAVER. I-Impute optimizes continuous similarities and dropout probabilities iteratively to reach a self-consistent state. However, these tools focus on imputation only. Directly applying the imputed expression profile to downstream tasks like cell population detection and cell-cycle inference might fail due to the curse of dimensionality (22, 23).

In previous work, we displayed matrix factorization-based model could conduct robust imputation ability alongside cancer subtype reversed dimension reduction in bulk cancer omics data (24). Thus, in this study, we developed SMURF to impute scRNA data based on matrix factorization (Fig.1). SMURF leverages a mixture of Poisson-Gamma distribution to deal with the zero-inflated problem. SMURF embeds cells and genes data into their latent space that facilities cell type or cell-type identification and visualization. We first accessed SMURF’s gene expression recovery ability with the state-of-the-art imputation tools I-Impute, SAVER, MAGIC, and kNN-smoothing. In *in silico* data assessment, SMURF models paraded the most robust gene expression recovery power with low RMSE and high Pearson correlation. Moreover, SMURF’s cell latent matrices (*H*) successfully preserved the underlying cell-subpopulation structures against full-dimensional imputed cell-gene expression data for different sparsity. As for the wet lab dataset evaluation, we demonstrated that SMURF recovered the genes’ distribution from WM989 Drop-seq data. We further collected eight wet lab cell line datasets to benchmark the tools. SMURF exhibited feasible cell subpopulation discovery efficacy on all the eight cell line mixtures. Furthermore, we illustrated that SMURF could embed the cells into a 1D-oval latent space and preserve the time course of the cell cycle.

**Fig. 1.**
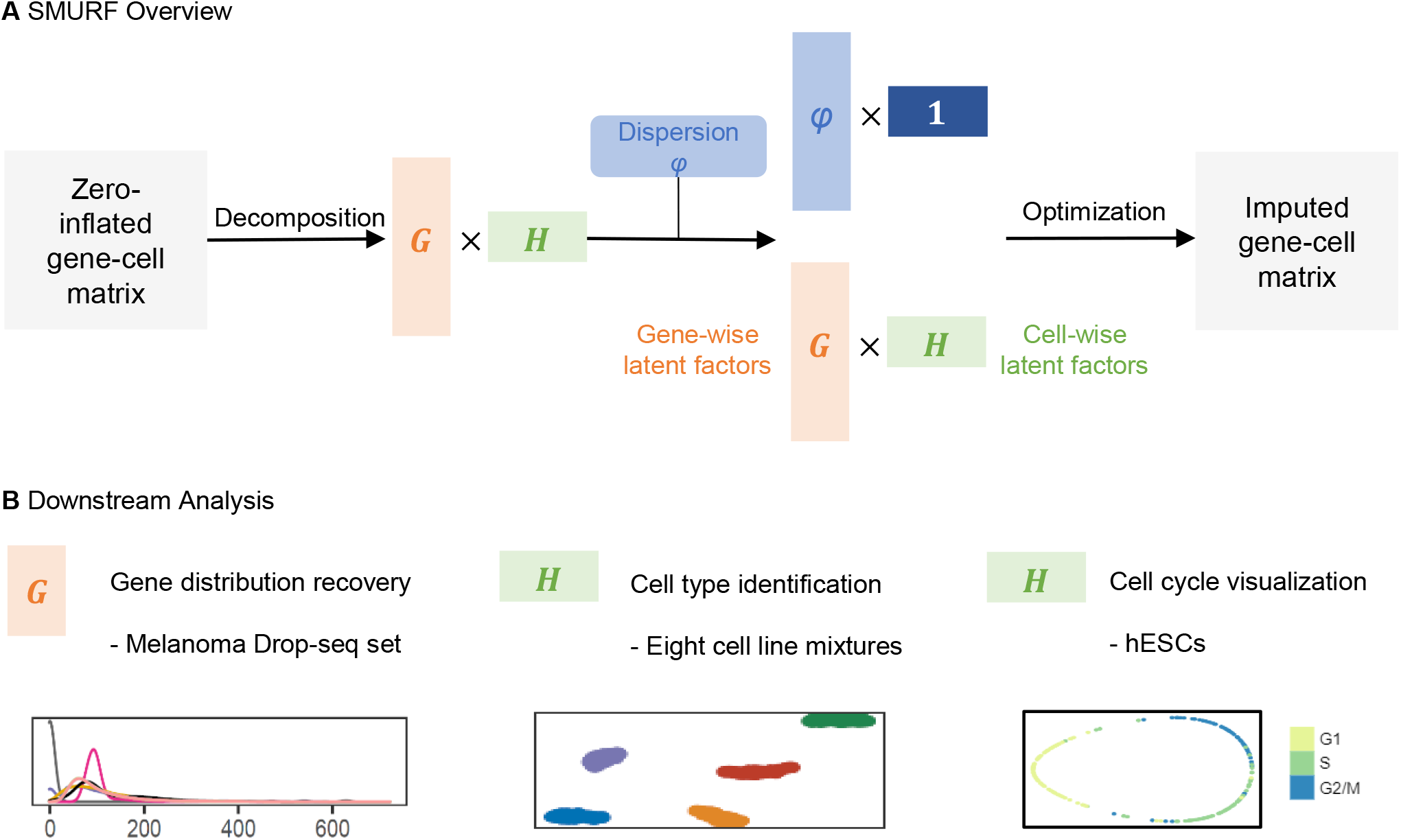
Overview of SMURF

## Methods

### Algorithm of SMURF

#### Problem formulation

Let ***X*** ∈ ℝ^*M* ×*N*^ be the normalized gene-cell matrix, where *M* and *N* is the number of genes and cells, respectively. Here we use a library size normalization defined as the library size divided by the mean library size across all cells, but SMURF can also be applied to prenormalized dataset using different normalization method. SMURF assumes that the element ***X***_*ij*_ in ***X*** follows a negative binomial distribution, which can be decomposed into a mixture of Poisson and Gamma distribution.

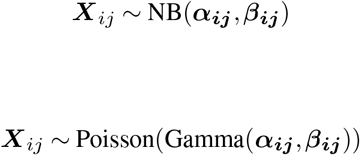

where ***α***_***ij***_ is the shape parameter and ***β***_***ij***_ is the scale parameter of Gamma distribution.These two parameters actually reflect the mean ***µ***_***ij***_ and variance ***ν***_***ij***_ of the distribution.

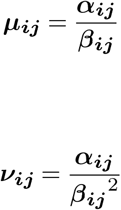

Our goal is to make a reasonable estimate of the prior mean ***µ*** and prior variance ***ν*** through matrix factorization based on the negative binomial distribution, and use the calculated prior mean ***µ*** as the final estimate of the gene expression level of this model (Algorithm 1).

##### Algorithm 1 SMURF

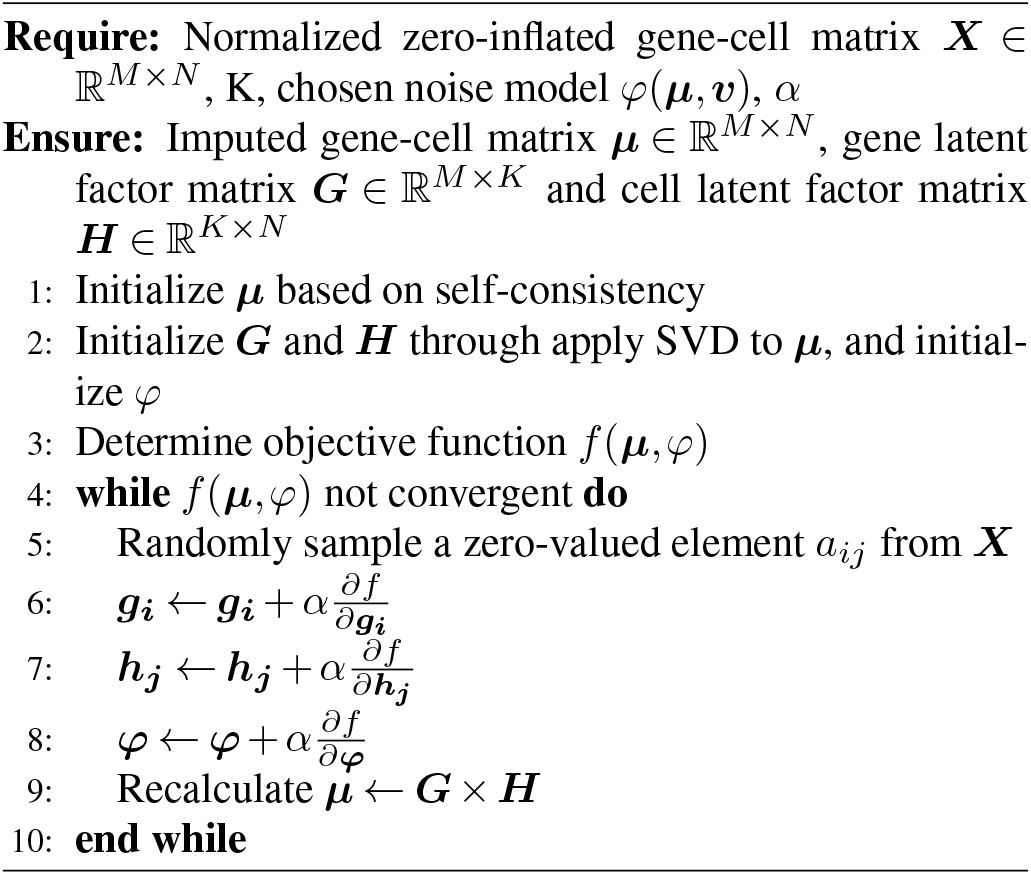

#### Model initialization based on self-consistency

In order to improve the accuracy of the model, it is necessary to initialize the parameters. The initialization process not only needs to determine the initial values of the parameters, but also needs to disassemble the parameters into two matrices to facilitate subsequent matrix decomposition.

First, we initialize the mean ***µ*** based on the self-consistency. As stated in I-Impute (21), a self-consistent functional mapping *f* : *x* → *x* given by input data ***X*** ∈ *M* × *N* is:

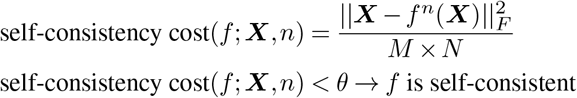

where *n* is the function iteration times and *θ* stands for the tolerant noise.

Here we assume that ***µ***_***ij***_ is a prediction derived by other genes in the same cell defined by the inter-gene relationship matrix ***R*** ∈ ℝ^*M* ×*M*^, where *r*_*ij*_ in ***R*** represents the relation coefficient of gene *j* for gene *i*. Obviously, *r*_*ij*_ = 0 when *i* = *j*. So ideally, the following equation should exist:

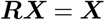

We call the above equation as the *first-order* equation, from which we can continue to obtain the following *high-order* equations:

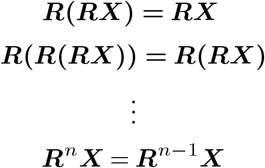

It is the inherent property of the relation matrix ***R*** to make the above first to *n*-order equations true. The *n*-order equations is the iterative function mapping *f* (***X***) = ***RX*** refining ***X*** to achieve self-consistency. Based on this property, we can calculate the approximate matrix 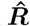 of the relationship matrix that satisfies the self-consistency nature, and make coarse imputation of the cell gene matrix to get the initialization of ***µ***.

In order to make the size of all parameter values in 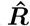 reasonable, and to enhance the stability and expressive ability of the model, we first initialize a matrix ***D*** ∈ 𝕄 × 𝕄, where

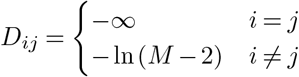

And we can initialize relation matrix 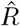 as *sigmoid*(*D*), where

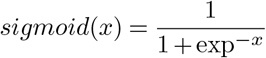

Then we design the objective function based on selfconsistency:

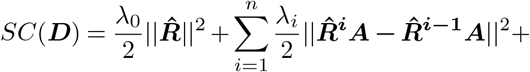

We can get a reasonable approximation of the relationship matrix by optimizing the above objective function. In practice, n does not need to be too large, generally 3 to 5 can be used to obtain satisfactory results.Finally, we can get a preliminary prediction of ***µ***:

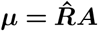

Above, we have applied the inter-gene relationship matrix for discussion. In fact, this idea is also applicable to the intercell relationship matrix. However, we have found in actual experiments that since the number of cells in the relevant data set is far less than the number of genes, when we initialize with the self-consistency of the inter-cell relationship matrix, although it consumes less computing resources, the case of underfitting occurs more often.

Based on singular value decomposition, we decompose the matrix ***µ*** into two matrices ***G*** and ***H***, where ***G*** ∈ ℝ^*M* ×*K*^, ***H*** ∈ ℝ^*K*×*N*^, *K* ≪ *M, N* .

For the prior variance ***ν***, we assume that for each gene, there is an implicit connection between its mean and variance, and introduce three different noise models to estimate the variance: constant variance model, constant Fano model, and constant coefficient of variation model, detail formulation is in Supplementary Methods.

#### Objectives

After initialization, SMURF uses the matrix decomposition based on the negative binomial distribution and different noise model to denoise the original expression matrix ***X*** and impute in the dropouts in the matrix. First, we need to obtain the set *P* composed of the coordinates of cells and genes according to the matrix ***X***, and make *P* satisfy ∀(*i, j*) ∈ *P*, ***X***_*ij*_ > 0.

For the constant variance model, the objective function is

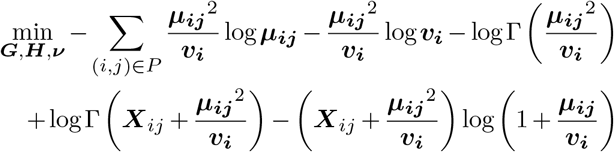

with ***µ*** = ***G*** × ***H***. Let the loss function be *f*_*cv*_ and calculate separately 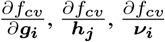, through the gradient descent method, continuously update the mean ***µ*** and variance ***ν*** until finally converges.

Under the assumption of the constant Fano model, the objective function is

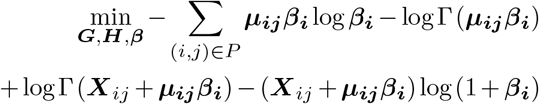

In practice, for numerical stability, we calculate 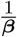 instead of ***β***

Similarly, for the constant coefficient of variance model, we can get the objective function

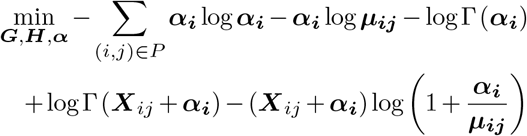

In practice, for numerical stability, we calculate 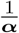 instead of ***α***.

### Simulation and benchmark settings

We used Splatter (25) to simulate five scRNA-seq datasets. The parameter configurations were nGenes=2000, batch-Cells=500, seeds=42, group.prob = c(0.2, 0.2, 0.2, 0.2, 0.2), dropout.type=“experiment”, dropout.shape=-1 and droupout.mid=1, 3, 4, 5, and 7 for dropout rate 28.38%, 63.14%, 78.00%, 88.06%, and 97.22%, respectively.

We selected I-Impute, SAVER, MAGIC, and kNN-smoothing as state-of-art imputation tools. For the SAVER R package, we used the “saver” function with the parameters ncores=12 and estimates.only=TRUE to perform the imputation tasks. The parameters for scImpute are drop_thre=0.5, ncores=10, Kclusters=(number of true clusters in input data). For I-Impute, we used n=20, c_drop=0.5, p_pca=0.4, normalize=False for synthetic data and True for real data sets, iteration=True. For MAGIC, genes=“all_genes”, knn=5, delay=1,t=3. The parameters for kNN-smoothing are k=10, d=10, dither=0.03

## Result

### SMURF recovers the dropout gene expression in *in silico* trials

We checked out whether SMURF can recover the dropout gene expression in *in silico* scRNA-seq data generated by Splatter (25). We simulated a raw read count matrix with 2,000 genes and five cell groups, each group with 100 cells. Next, we replaced 28.38%, 63.14%, 78.00%, 88.06%, and 97.22% of non-zero values with zero, yielding five zero-inflated matrices with sparsity of 42.86%, 70.59%, 82.45%, 90.47%, 97.79%.

We evaluated the consistency of imputed values with the raw ones using root mean square error (RMSE) and Pearson correlation (Cor.). kNN-smoothing exhibited high RMSE errors while I-Impute and three SMURF variance modes (CV, F, and V) had low RMSE. SMURF achieved the lowest RMSE when the sparsity reaches 80% (Fig.2A). Then, we measured the Pearson correlation between raw and recovered matrix entries (Fig.2A and Supplementary Fig.S1). For 28.38% dropout rate, all tools achieved (Cor. > 0.92, *p <* 2.2e-16)). The correlation scatterplots demonstrate MAGIC and SMURF models outperformed other tools when sparsity goes up with no severe drop-down or pull-up estimation. SMURF models had slightly larger correlations than MAGIC. For 63.14% dropout rate, MAGIC, SMURF_CV, SMURF_F, and SMURF_V had Cor. value of 0.926, 0.933, 0.9323, 0.933, respectively (*p <* 2.2e-16). For 78.00% dropout rate, MAGIC, SMURF_CV, SMURF_F, and SMURF_V had Cor. value of 0.900, 0.913, 0.913, 0.913, respectively (*p <* 2.2e-16). For 88.06% dropout rate, MAGIC, SMURF_CV, SMURF_F, and SMURF_V had Cor. value of 0.856, 0.873, 0.874, 0.873, respectively (*p <* 2.2e-16). For 97.22% dropout rate, MAGIC, SMURF_CV, SMURF_F, and SMURF_V had Cor. value of 0.727, 0.746, 0.745, 0.744, respectively (*p <* 2.2e-16). I-Impute, kNN-smoothing, and SAVER tend to pull down some highly expressed genes, hence producing relatively low correlation values. Furthermore, we inspected the group marker genes heatmaps using raw, dropout, and recovered expression levels (Supplementary Fig.S2). MAGIC and SMURF’s marker gene matrices are closest to the ground truth. On 78.00% and 88.06% dropout sets, I-Impute, SAVER, and kNN-smoothing failed to reproduce a clear marker expression pattern. However, all tools were unable to recover the marker expression pattern when the dropout rate reaches 97.22%.

**Fig. 2.**
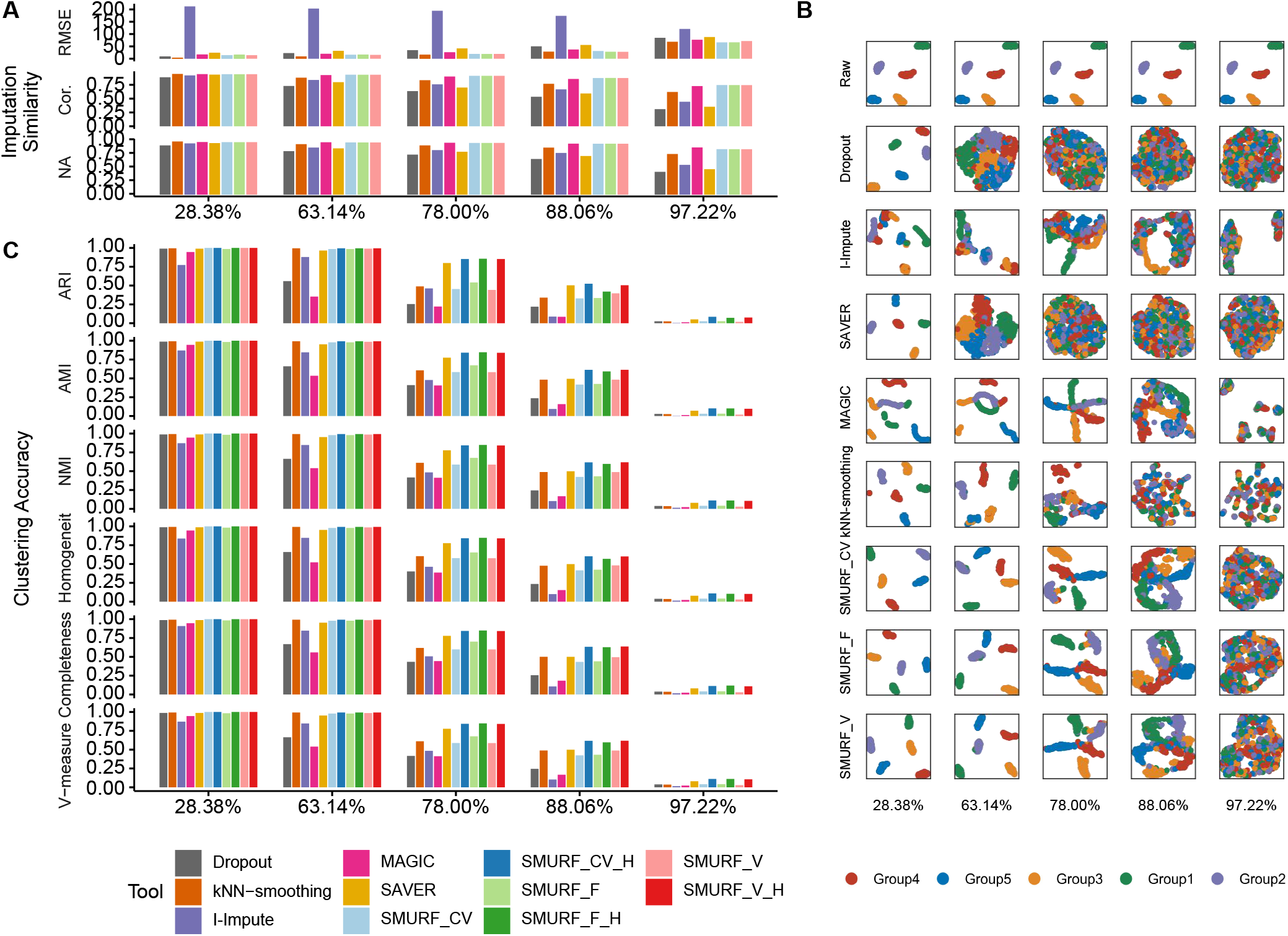
Benchmarking result on five *in silico* scRNA data with dropout rate of 28.38%, 63.14%, 78.00%, 88.06%, and 97.22%. (A) The imputation consistency was measured by root mean square error (RMSE) and Pearson correlation (Cor.). (B) UMAP plot. Cells from the same group share the same annotation color. From left to right are datasets with the dropout rate of 28.38%, 63.14%, 78.00%, 88.06%, and 97.22%. From top and bottom are tools: raw, dropout, I-Impute, SAVER, MAGIC, kNN-smoothing, SMURF_CV, SMURF_F, and SMURF_V. (C) The clustering accuracy was measured by adjusted rand index (ARI), adjusted mutual information (AMI), normalized mutual information (NMI), homogeneity score (Homogeneity), completeness score (Completeness), and V-measure score (V-measure). SMURF_CV, SMURF_V, and SMURF_F stand for SMURF with variance mode constant coefficient of variance (CV), constant Fano (F), and constant variance (V), respectively. SMURF models with suffix “H” mean the clustering was conducted with the cell latent factors yielded by SMURF.

We then assessed whether SMURF maintains the cell subgroups structures well while retrieving the missing entries. Fig.2B displays the UMAP embedding plots of the raw, dropout, and recovered matrices annotated with five cell groups. The UMAP plot of expression data recovered by MAGIC and SMURF separated the five cell groups even when the dropout rate reached 88.06%. The UMAP plot of kNN-smoothing blurred at the dropout rate of 78%. I-Impute and SAVER performance worst, failing at 63.14%. Furthermore, we employed spectral clustering to the dropout, recovered, and SMURF’s cell latent (*H*) matrices. We evaluated the clustering accuracy using six metrics: adjusted Rand index (ARI) (26), adjusted mutual information (AMI) (27), normalized mutual information (NMI) (28), homogeneity score (Homogeneity) (29), completeness score (Completeness) (29), and V-measure score (V-measure) (29). These metrics measure the degree of overlap between the estimation and ground truth clusters. The value of 0 signifies random clustering while 1 indicates perfectly complete labeling. In Fig.2C, the overall clustering accuracy declined with the dropout rate increased. kNN-smoothing and MAGIC exhibited the lowest clustering accuracy in all settings, sometimes even worse than the dropout case. The expression data recovered by I-Impute, SAVER, SMURF_CV, SMURF_F, SMURF_V displayed moderate improvements on clustering accuracy against the dropout data. Notably, the clustering accuracy yielded by the cell latent matrices *H* outperformed all high dimensional cell expression data and manifested the most desirable cell group classification power. Especially, SMURF_CV_H, SMURF_F_H, SMURF_V_H obtained the most noticeable clustering accuracy enhancements against their corresponding full matrices (SMURF_CV, SMURF_F, SMURF_V) in 78%, 88.06%, and 97.22% datasets.

Overall, the *in silico* assessment indicated that SMURF models paraded the most robust gene expression recovery power. Moreover, SMURF’s cell latent matrices *H* successfully preserved the underlying cell-subpopulation structures.

### SMURF enable accurate unsupervised clustering on mixture of cell line data

We applied SMURF to eight wet-lab mixture cell line datasets to benchmark the cell sub-population problem. The first set is p3cl, including human dermal fibroblasts-skin, breast cancer luminal epithelial cell line MCF-7, and breast cancer basal-like epithelial cell line MDA-MB-468 in a 1:3:6 ratio (30, 31). We evaluated SMURF with CellBench (32), where five human lung adenocarcinoma cell lines were mixed and sequenced with 10x Genomics, CEL-seq2, and Drop-seq, yielding seven datasets (mixture of HCC827, H1975, and H228: sc_10x, sc_celseq2, sc_dropseq; mixture of HCC827, H1975, A549, H838, and H2228: sc_10x_5cl, sc_celseq2_5cl_p1, sc_celseq2_5cl_p2, sc_celseq2_5cl_p3).

Likewise, we evaluated the consistency of imputed values with the raw ones using RMSE and Pearson correlation. kNN-smoothing exhibited high RMSE errors while SMURF and other tools had relatively low RMSE (Fig.3A). MAGIC had the highest Pearson correlation between raw and recovered matrix entries (Fig.3A). The performance of I-Impute and SAVER performance is similar to that of dropout data. The SMURF models has the second place in five out of eight sets (sc_celseq2_5cl_p1, sc_celseq2_5cl_p2, sc_celseq2_5cl_p3, sc_celseq2, and sc_dropseq). The investigation of cell type marker genes heatmaps using raw, dropout, and recovered expression levels (Supplementary Fig. S3) demonstrates that the SMURF’s marker gene matrices are closest to the raw data on all eight sets. All other tools failed to produced distinct cell type pattern in some datasets. We then evaluated whether SMURF keeps the cell type structures well while retrieving the missing entries. Fig.3B illustrates the UMAP embedding plots of the raw, dropout, and recovered matrices annotated with the cell line labels. The UMAP plot of expression data recovered by I-Impute, SAVER and SMURF drew distinct cell line groups. However, MAGIC and kNN-smoothing mixed different cell line together in a mess. Furthermore, Fig.3C further suggested that MAGIC and kNN-smoothing had the poorest spectral clustering accuracy on 600 of highly variable genes (HVG).

**Fig. 3.**
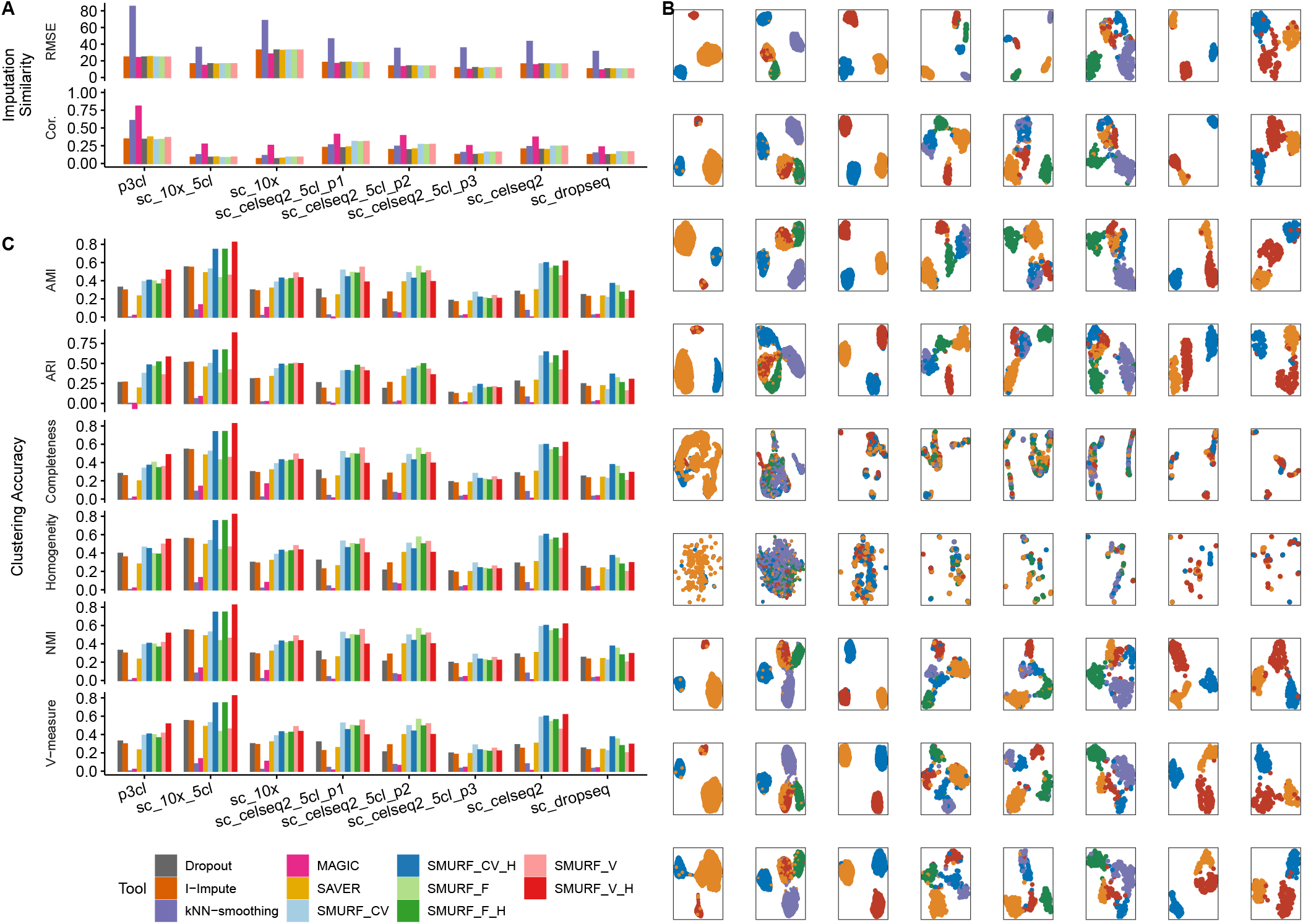
Benchmarking result on eight mixture of cell line scRNA data. (A) The imputation consistency was measured by root mean square error (RMSE) and Pearson correlation (Cor.). (B) UMAP plot. Cells from the same group share the same annotation color. From left to right are datasets p3cl, sc_10x_5cl, sc_10x, sc_celseq2_5cl_p1, sc_celseq2_5cl_p2, sc_celseq2_5cl_p3, sc_celseq2, and sc_dropseq with the dropout rate of 90%, 80%, 80%, 65%, 65%, 75%, 65%, 75% respectively. From top and bottom are tools: Raw, Dropout, I-Impute, SAVER, MAGIC, kNN-smoothing, SMURF_CV, SMURF_F, and SMURF_V. (C) The clustering accuracy was measured by adjusted rand index (ARI), adjusted mutual information (AMI), normalized mutual information (NMI), homogeneity score (Homogeneity), completeness score (Completeness), and V-measure score (V-measure) of HVG600. SMURF_CV, SMURF_V, and SMURF_F stand for SMURF with variance mode constant coefficient of variance (CV), constant Fano (F), and constant variance (V), respectively. SMURF models with suffix “H” mean the clustering was conducted with the cell latent factors yielded by SMURF.

Overall, kNN-smoothing and MAGIC demonstrated the lowest clustering accuracy in all cell line mixtures, even worse than the dropout case. The expression data recovered by I-Impute and SAVER displayed moderate clustering accuracy improvements against the dropout on sc_celseq2_5cl_p2. Notably, the clustering accuracy yielded by SMURF outperformed on all cell mixtures and displayed the most desirable cell group classification power. Especially, cell latent matrices *H* obtained the most noticeable clustering accuracy enhancements against all corresponding full matrices in the majority of datasets.

In all, the cell line mixture assessment indicated that SMURF models paraded the most robust gene expression recovery power. Moreover, SMURF’s cell latent matrices *H* successfully preserved the underlying cell-subpopulation structures.

### SMURF facilitates pesudo-time prediction for cell– cycle data

We leveraged six cell-cycle datasets to show SMURF facilitates pesudo-time prediction. In H1-hESC datasets, 247 cells were identified as being in G0/G1, S, or G2/M phases using fluorescent ubiquitination-based cell-cycle indicators (33). With G0/G1, S, and G2/M phases labeled by Hoechst staining, mESC-Quartz and mESC-SMARTer has 23 and 288 mESCs sequenced by Quartz-seq and SMARTer, respectively (34, 35). Then, as the 3Line-qPCR dataset provides G0/G1, S, and G2/M cell stages marked by Hoechst staining as well, we split it into three datasets 3Line-qPCR_H9 (227 cells), 3Line-qPCR_MB (342 cells), and 3Line-qPCR_PC3 (361 cells) as the pesudo-time prediction benchmark sets (36).

Likewise, we first adopted I-Impute, kNN-smoothing, MAGIC, SAVER, and SMURF to impute the six datasets. Then, we used SMURF to embed the raw and imputed full-dimensional expression profile and *H* into 1D-oval latent space (Fig.4B). In the 1D-oval plot of raw H1-hESC, the cells from the G2/M phase are split into two subgroups, and the cells are not ordered according to the correct time course. However, in the 1D-oval plot of SMURF’s cell latent matrix *H*, cells are formed a G0/G1-S-G2/M cycle alongside the oval. We further adopted change index (CI) and Pearson correlation coefficient (PCC) (5) to quantitatively measure the accuracy of 1D-oval pseudo-time order against known cell cycle phases labels. A close to 1 value of CI and PCC signifies near perfect time-series order, while a value of 0 means the predict psueo-time order is randomly matched with the ground-truth cell-cycle order. The SMURF_H has higher CI and PCC than SMURF and other imputation tools at most cases. This signifies SMURF can reduce the cell-cycle noise utilizing matrix factorization, thus facilitating the cell-cycle pseudo-time prediction.

**Fig. 4.**
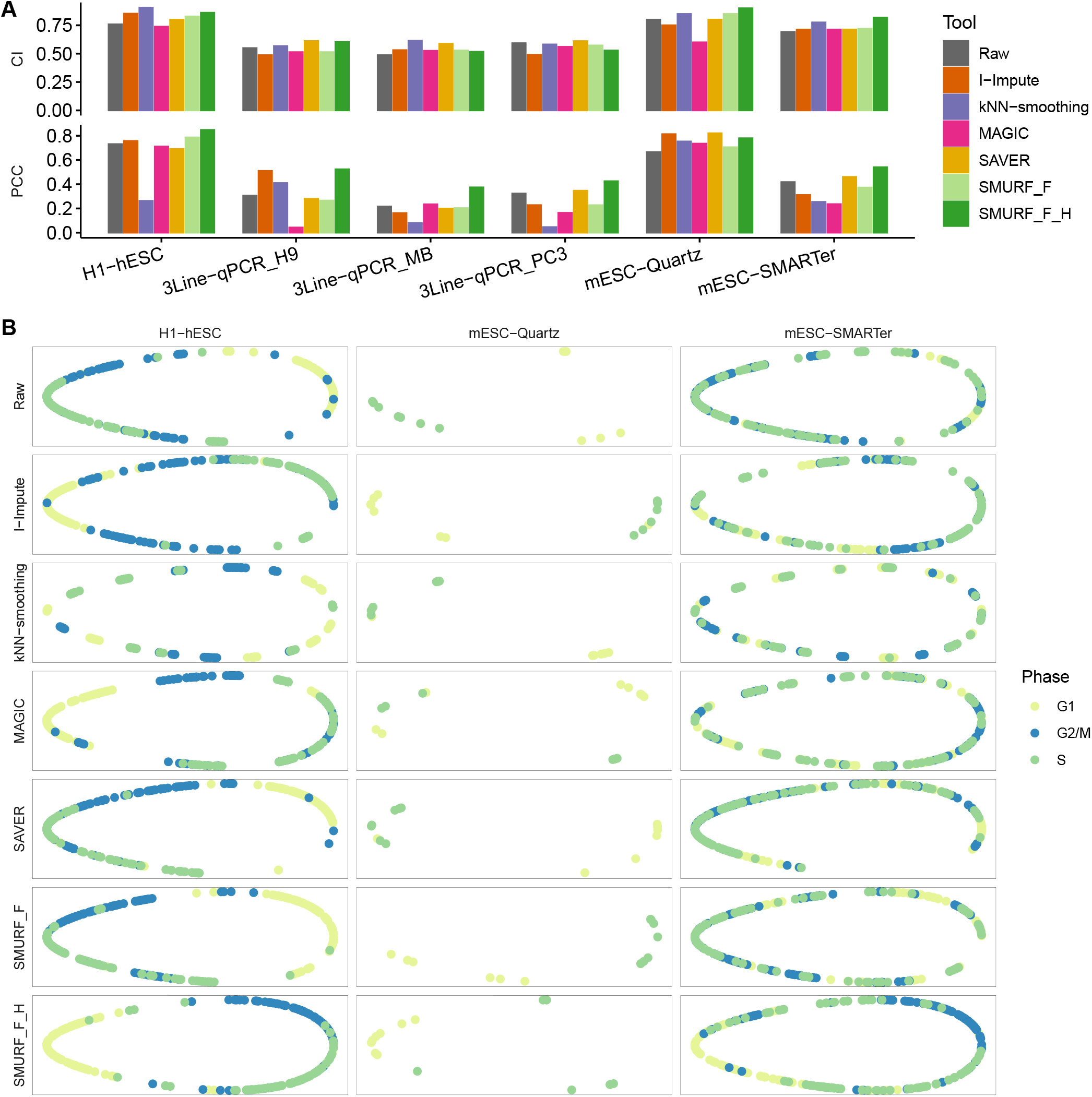
Cell-cycle visualization for hESCs. (A) The time series scoring accuracy measured by the change index (CI) and Pearson correlation coefficient (PCC). (B) 1D oval plot. Cells from the same cell cycle phase share the same annotation color. From left to right are datasets: H1-hESC, mESC-Quartz, and mESC-SMARTer. From top and bottom are tools: Raw, I-Impute, kNN-smoothing, MAGIC, SAVER, SMURF_F, and SMURF_F_H.

### SMURF recovers the distribution of genes from WM989 Drop-seq data

We benchmarked SMURF with the melanoma cell line data, WM989, where Torre *et al*. profiled 8,498 WM989 single cells employing Drop-seq (37) and Snaffer *et al*. measured the golden-standard expression of 26 drug-resistant marker and housekeeping genes among 7,000–88,000 WM989 cells with single-molecule RNA fluorescence *in situ* hybridization (smRNA FISH) (38). As 15 genes overlapped between and the sequenced cells are different from the two approaches (37, 38). We assessed imputation performance by comparing these 15 gene distributions and correlations of imputation tools recovered to distributions captured by gold-standard smRNA FISH (Fig.5). I-Impute is not compared here as it failed finished the task in 24 hours.

**Fig. 5.**
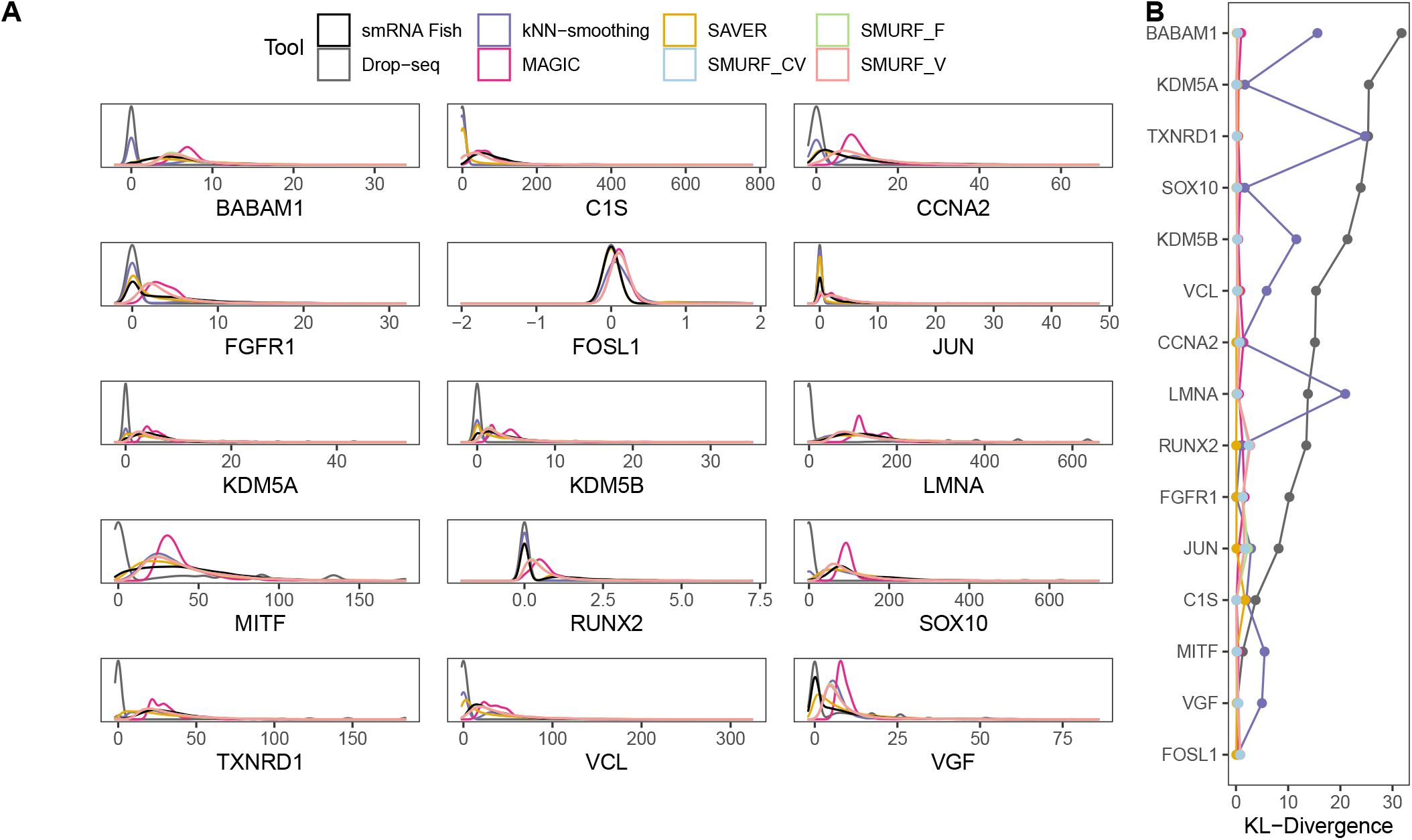
Benchmarking result on melanoma cell line WM989 (A) The imputed gene distribution and FISH estimated gene distribution among 15 genes. (B) The Kullback–Leibler (KL) divergence between imputed gene distribution and FISH estimated gene distribution.

We drew the density plot of each gene’s expression across cells (Fig.5A) and used Kullback–Leibler (KL) divergence (39) to measure how gene distributions recovered by imputation tools were different from the reference gene distributions estimated by smRNA FISH (Fig.5B). Compared with Drop-seq, the expression distributions obtained by SMURF and SAVER were closer to FISH. MAGIC and kNN-smoothing demonstrated a worse imputation ability as MAGIC on the shape of density plot and kNN-smoothing show the largest KL divergence on several genes.

## Discussion

In this study, we presented SMURF to impute scRNA data leveraging a mixture of Poisson-Gamma distribution and embed cells and genes data into their latent space with matrix factorization. Experiments using *in silico* and wet lab data illustrated the imputation and dimension reduction efficacy. We first accessed SMURF’s gene expression recovery ability with the state-of-the-art imputation tools I-Impute, SAVER, MAGIC, and kNN-smoothing. In synthetic data assessment, SMURF models paraded the most robust gene expression recovery power with low RMSE and high Pearson correlation. SMURF successfully recovered the gene distribution from melanoma cell line Drop-seq data. Moreover, we demonstrated SMURF’s cell latent matrices (*H*) successfully preserved the underlying cell-subpopulation structures for different sparsity against full dimensional imputed cell-gene expression data on synthetic datasets and eight cell line mixtures datasets. As for the wet lab dataset evaluation, we demonstrated that SMURF recovered the genes’ distribution from WM989 Drop-seq data. We further collected eight wet lab cell line datasets to benchmark the tools. SMURF exhibited feasible cell subpopulation discovery efficacy on all the eight cell line mixtures. Furthermore, we illustrated that SMURF could embed the cells into a 1D-oval latent space and preserve the time course of the cell cycle.

Both SMURF and SAVER adopt Poisson-Gamma distribution to the zero-inflated fact of single-cell data. The advantage of SMURF against SAVER and other imputation tools is the support dimensional reduction utilization matrix factorization. We model the distribution mean ***µ***_*ij*_ with the dot product of two *K*-dimensional vectors ***g***_***i***_ and ***h***_***j***_, which signifies the low-dimensional representation of gene and cell in the latent space, respectively. The latent factor ***g*** and ***h*** preserve the underlying gene-gene interactions and cell-cell structures and overcome the curse of dimensionality. Thus, the latent matrix *H* manifested better cell type detection power against the full-dimensional matrix. Furthermore, SMURF provides 1D-oval manifold embedding dedicated for cell-cycle visualization.

However, the selection of the reduced dimension *K* is still an open question. Generally, the value of K first depends on the size of the zero-inflated matrix. The number of cells or genes should be at least several times larger than K, otherwise the model may be over-fitting and not effective. The dropout rate in the matrix also affects the value of K. For a matrix with a small dropout rate, the value of K can be increased appropriately, while for a very sparse matrix, a smaller K should be tried.

## Data availability

The p3cl was obtained from (30, 31). The 3Line-qPCR_PC3 was obtained from (36). The CellBench data was obtained from (32). The H1-hESC data was obtained from (33). The mESC-Quartz was obtained from (34). The mESC-SMARTer was obtained from (35). The WM989 Drop-seq was obtained with accession code GSE99330 (37). The WM989 smRNA FISH data was obtained from the link https://www.dropbox.com/s/ia9x0iom6dwueix/fishSubset.txt?dl=0 (38).

## Software availability

The source code of SMURF is available at https://github.com/deepomicslab/SMURF.

## Competing interests

The authors declare that they have no competing interests.

## Acknowledgments

Not applicable.

## Funding

This project was supported by SIRG (CityU SIRG 7020005).

## Supplementary Note 1: Supplementary Method

### 1.1. Variance model

The first model we introduced is the constant variance model. We consider the scenario ***ν***_***ij***_ = ***ν***_***i***_, all cells share the same constant dispersion factor for the specified gene. Therefore, the negative binomial distribution has the following form in this case:

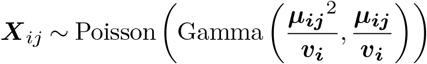

From this we can get the probability density function of the negative binomial distribution and initialize the prior variance through maximum likelihood estimation.

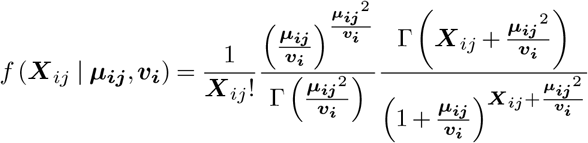

For all elements belong to *P*, we can multiply the density functions to get the likelihood function.

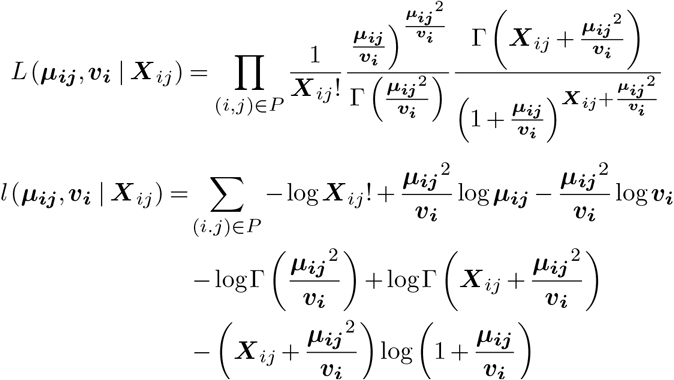

The second model we introduced is the constant Fano model, which assumes ***ν***_***ij***_ = ***F***_***i***_***µ***_***ij***_, the Fano Factor can be expressed as

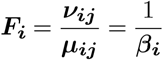

Thus, we have

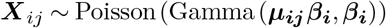

Here we estimate ***β*** instead of directly estimating ***ν***, and calculate the likelihood in a similar way as above

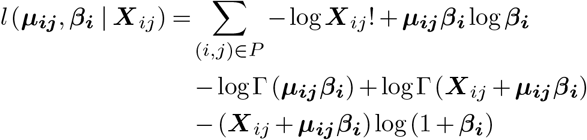

The last model introduced is constant coefficient of variation model. Under the assumptions of this model, there is a quadratic relationship between the mean and the variance, which is ***ν***_***ij***_ = *C****V***_***i***_^2^***µ***_***ij***_^2^, and we have

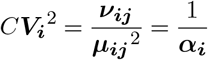

Similar to the Fano model, we can get the following log-likelihood

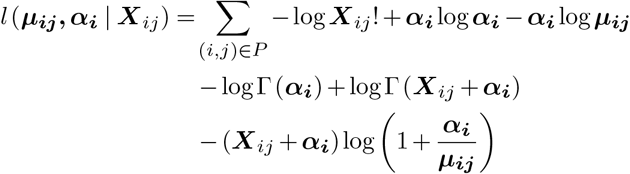

### Supplementary Note 2: Supplementary File

**Fig. S1.**
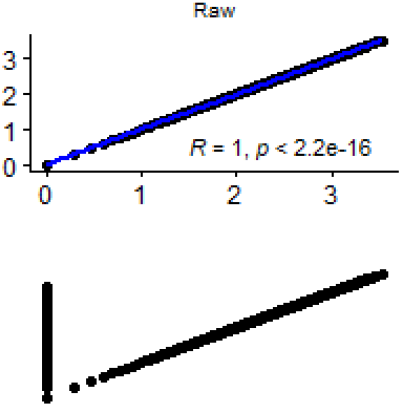
Scatter plot and Pearson correlation of gene expression in cells between ground-truth and specified tools, (X-axis: log10(count + 1) in Raw; Y-axis: log10(count + 1) in Tool). From left to right are datasets with the dropout rate of 28.38%, 63.14%, 78.00%, 88.06%, and 97.22%. From top and bottom are tools: raw, dropout, I-Impute, SAVER, MAGIC, kNN-smoothing, SMURF_CV, SMURF_F, and SMURF_V. SMURF_CV, SMURF_V, and SMURF_F stand for SMURF with variance mode constant coefficient of variance (CV), constant Fano (F), and constant variance (V), respectively.

**Fig. S2.**
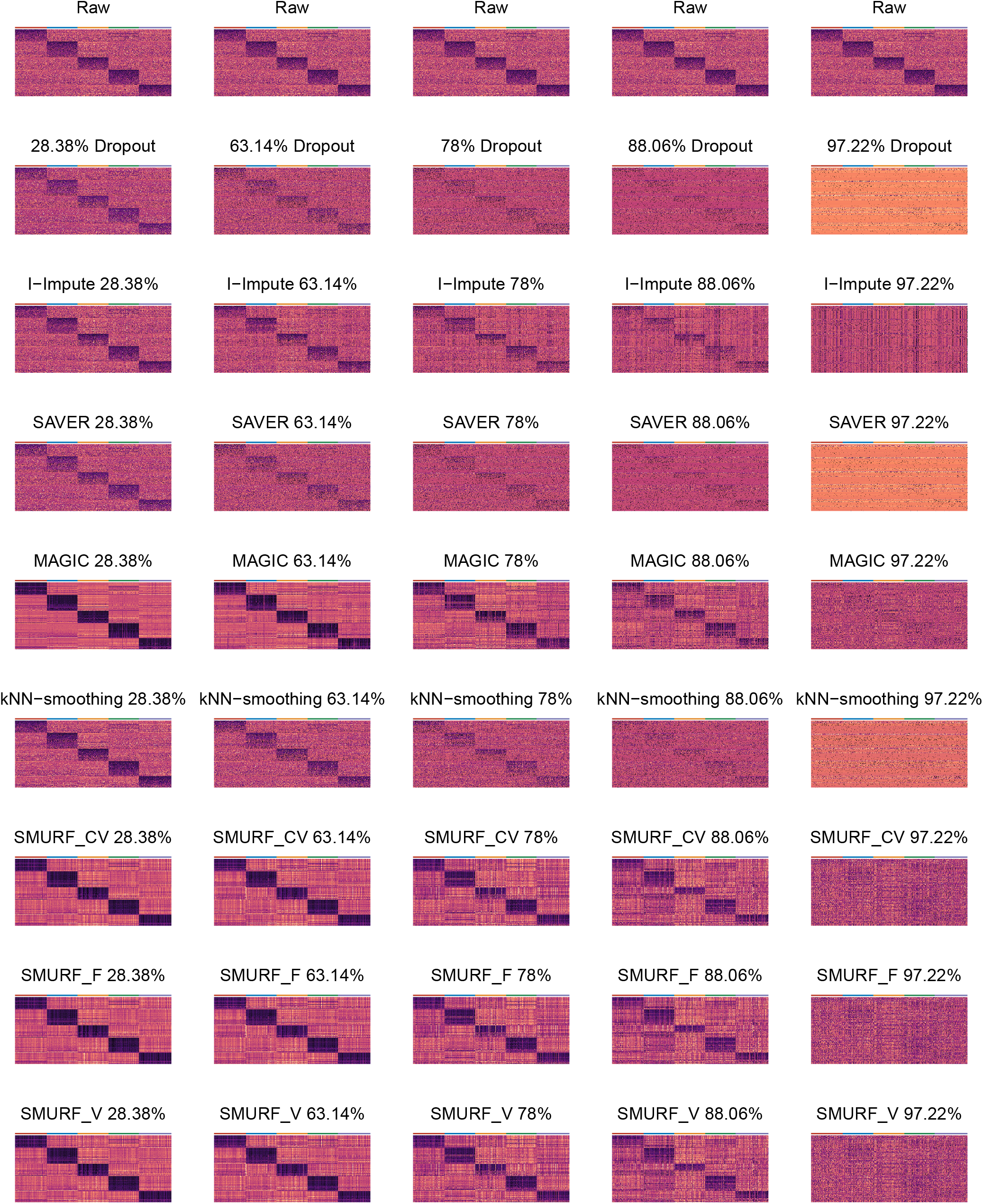
Marker gene expression heatmap with columns as genes and rows as cells. Cells from the same group share the same annotation color. From left to right are datasets with the dropout rate of 28.38%, 63.14%, 78.00%, 88.06%, and 97.22%. From top and bottom are tools: raw, dropout, I-Impute, SAVER, MAGIC, kNN-smoothing, SMURF_CV, SMURF_F, and SMURF_V. SMURF_CV, SMURF_V, and SMURF_F stand for SMURF with variance mode constant coefficient of variance (CV), constant Fano (F), and constant variance (V), respectively.

**Fig. S3.**
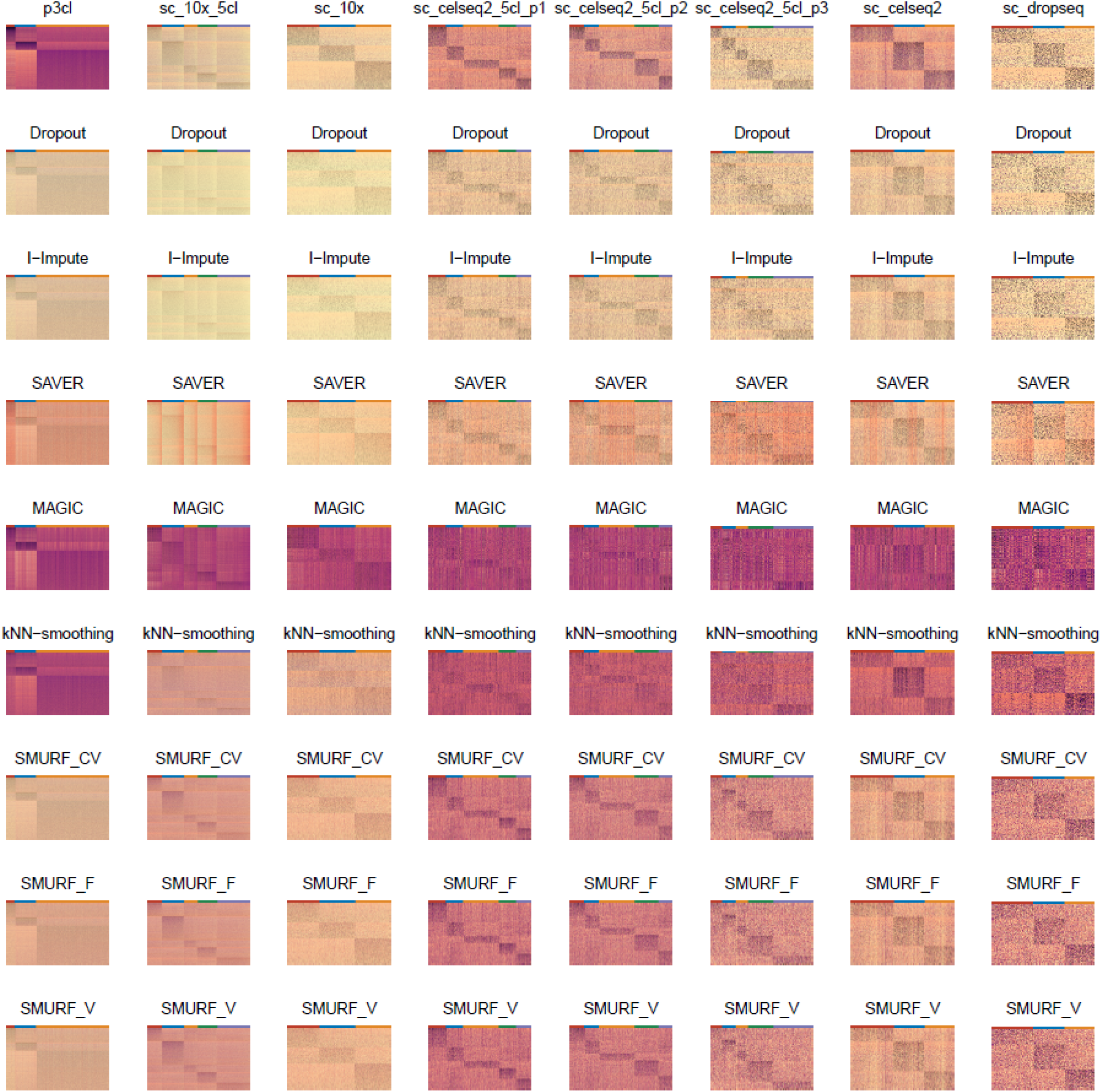
Marker gene expression heatmap with columns as genes and rows as cells. Cells from the same group share the same annotation color. From left to right are datasets p3cl, sc_10x_5cl, sc_10x, sc_celseq2_5cl, sc_celseq2_5cl_p1, sc_celseq2_5cl_p2, sc_celseq2_5cl_p3, sc_celseq2, and sc_dropseq. From top and bottom are tools: raw, dropout, I-Impute, SAVER, MAGIC, kNN-smoothing, SMURF_CV, SMURF_F, and SMURF_V. SMURF_CV, SMURF_V, and SMURF_F stand for SMURF with variance mode constant coefficient of variance (CV), constant Fano (F), and constant variance (V), respectively.

## Notes

### Competing Interest Statement

The authors have declared no competing interest.

## Reference

1. Antoine-Emmanuel Saliba, Alexander J Westermann, Stanislaw A Gorski, and Jörg Vogel. Single-cell rna-seq: advances and future challenges. Nucleic acids research, 42(14):8845–8860, 2014.

2. Aleksandra A Kolodziejczyk, Jong Kyoung Kim, Valentine Svensson, John C Marioni, and Sarah A Teichmann. The technology and biology of single-cell rna sequencing. Molecular </p> cell, 58(4):610–620, 2015.

3. Serena Liu and Cole Trapnell. Single-cell transcriptome sequencing: recent advances and remaining challenges. F1000Research, 5, 2016.

4. Cole Trapnell, Davide Cacchiarelli, Jonna Grimsby, Prapti Pokharel, Shuqiang Li, Michael Morse, Niall J Lennon, Kenneth J Livak, Tarjei S Mikkelsen, and John L Rinn. The dynamics and regulators of cell fate decisions are revealed by pseudotemporal ordering of single cells. Nature biotechnology, 32(4):381, 2014.

5. Zehua Liu, Huazhe Lou, Kaikun Xie, Hao Wang, Ning Chen, Oscar M Aparicio, Michael Q Zhang, Rui Jiang, and Ting Chen. Reconstructing cell cycle pseudo time-series via single-cell transcriptome data. Nature communications, 8(1):22, 2017.

6. Aaron M Horning, Yao Wang, Che-Kuang Lin, Anna D Louie, Rohit R Jadhav, Chia-Nung Hung, Chiou-Miin Wang, Chun-Lin Lin, Nameer B Kirma, Michael A Liss, et al. Single-cell rna-seq reveals a subpopulation of prostate cancer cells with enhanced cell-cycle–related transcription and attenuated androgen response. Cancer research, 78(4):853–864, 2018.

7. Kuti Baruch, Aleksandra Deczkowska, Neta Rosenzweig, Afroditi Tsitsou-Kampeli, Alaa Mohammad Sharif, Orit Matcovitch-Natan, Alexander Kertser, Eyal David, Ido Amit, and Michal Schwartz. Pd-1 immune checkpoint blockade reduces pathology and improves memory in mouse models of alzheimer’s disease. Nature medicine, 22(2):135, 2016.

8. Åsa Segerstolpe, Athanasia Palasantza, Pernilla Eliasson, Eva-Marie Andersson, Anne-Christine Andréasson, Xiaoyan Sun, Simone Picelli, Alan Sabirsh, Maryam Clausen, Magnus K Bjursell, et al. Single-cell transcriptome profiling of human pancreatic islets in health and type 2 diabetes. Cell metabolism, 24(4):593–607, 2016.

9. Nathan Lawlor, Joshy George, Mohan Bolisetty, Romy Kursawe, Lili Sun, V Sivakamasundari, Ina Kycia, Paul Robson, and Michael L Stitzel. Single-cell transcriptomes identify human islet cell signatures and reveal cell-type–specific expression changes in type 2 diabetes. Genome research, 27(2):208–222, 2017.

10. Woosung Chung, Hye Hyeon Eum, Hae-Ock Lee, Kyung-Min Lee, Han-Byoel Lee, Kyu-Tae Kim, Han Suk Ryu, Sangmin Kim, Jeong Eon Lee, Yeon Hee Park, et al. Single-cell rna-seq enables comprehensive tumour and immune cell profiling in primary breast cancer. Nature communications, 8:15081, 2017.

11. Mihriban Karaayvaz, Simona Cristea, Shawn M Gillespie, Anoop P Patel, Ravindra Mylvaganam, Christina C Luo, Michelle C Specht, Bradley E Bernstein, Franziska Michor, and Leif W Ellisen. Unravelling subclonal heterogeneity and aggressive disease states in tnbc through single-cell rna-seq. Nature communications, 9(1):3588, 2018.

12. Xinyi Guo, Yuanyuan Zhang, Liangtao Zheng, Chunhong Zheng, Jintao Song, Qiming Zhang, Boxi Kang, Zhouzerui Liu, Liang Jin, Rui Xing, et al. Global characterization of t cells in non-small-cell lung cancer by single-cell sequencing. Nature medicine, 24(7):978, 2018.

13. Charissa Kim, Ruli Gao, Emi Sei, Rachel Brandt, Johan Hartman, Thomas Hatschek, Nicola Crosetto, Theodoros Foukakis, and Nicholas E Navin. Chemoresistance evolution in triplenegative breast cancer delineated by single-cell sequencing. Cell, 173(4):879–893, 2018.

14. Michael Bartoschek, Nikolay Oskolkov, Matteo Bocci, John Lövrot, Christer Larsson, Mikael Sommarin, Chris D Madsen, David Lindgren, Gyula Pekar, Göran Karlsson, et al. Spatially and functionally distinct subclasses of breast cancer-associated fibroblasts revealed by single cell rna sequencing. Nature communications, 9(1):5150, 2018.

15. Peter V Kharchenko, Lev Silberstein, and David T Scadden. Bayesian approach to single-cell differential expression analysis. Nature methods, 11(7):740, 2014.

16. Wei Vivian Li and Jingyi Jessica Li. An accurate and robust imputation method scimpute for single-cell rna-seq data. Nature communications, 9(1):997, 2018.

17. David Van Dijk, Roshan Sharma, Juozas Nainys, Kristina Yim, Pooja Kathail, Ambrose J Carr, Cassandra Burdziak, Kevin R Moon, Christine L Chaffer, Diwakar Pattabiraman, et al. Recovering gene interactions from single-cell data using data diffusion. Cell, 174(3):716–729, 2018.

18. Florian Wagner, Yun Yan, and Itai Yanai. K-nearest neighbor smoothing for high-throughput single-cell rna-seq data. BioRxiv, page 217737, 2017.

19. Mo Huang, Jingshu Wang, Eduardo Torre, Hannah Dueck, Sydney Shaffer, Roberto Bonasio, John I Murray, Arjun Raj, Mingyao Li, and Nancy R Zhang. Saver: gene expression recovery for single-cell rna sequencing. Nature methods, 15(7):539, 2018.

20. Wenpin Hou, Zhicheng Ji, Hongkai Ji, and Stephanie C Hicks. A systematic evaluation of single-cell rna-sequencing imputation methods. Genome biology, 21(1):1–30, 2020.

21. Xikang Feng, Lingxi Chen, Zishuai Wang, and Shuai Cheng Li. I-impute: a self-consistent method to impute single cell rna sequencing data. BMC genomics, 21(10):1–9, 2020.

22. Tallulah S Andrews, Vladimir Yu Kiselev, Davis McCarthy, and Martin Hemberg. Tutorial: guidelines for the computational analysis of single-cell rna sequencing data. Nature protocols, 16(1):1–9, 2021.

23. Richa Nayak and Yasha Hasija. A hitchhiker’s guide to single-cell transcriptomics and data analysis pipelines. Genomics, 2021.

24. Lingxi Chen, Jiao Xu, and Shuai Cheng Li. Deepmf: deciphering the latent patterns in omics profiles with a deep learning method. BMC bioinformatics, 20(23):1–13, 2019.

25. Luke Zappia, Belinda Phipson, and Alicia Oshlack. Splatter: simulation of single-cell rna sequencing data. Genome biology, 18(1):174, 2017.

26. William M Rand. Objective criteria for the evaluation of clustering methods. Journal of the American Statistical association, 66(336):846–850, 1971.

27. Nguyen Xuan Vinh, Julien Epps, and James Bailey. Information theoretic measures for clusterings comparison: Variants, properties, normalization and correction for chance. The Journal of Machine Learning Research, 11:2837–2854, 2010.

28. Thomas M Cover and Joy A Thomas. Elements of information theory. John Wiley & Sons, 2012.

29. Andrew Rosenberg and Julia Hirschberg. V-measure: A conditional entropy-based external cluster evaluation measure. In Proceedings of the 2007 joint conference on empirical methods in natural language processing and computational natural language learning (EMNLP-CoNLL), pages 410–420, 2007.

30. Meichen Dong, Aatish Thennavan, Eugene Urrutia, Yun Li, Charles M Perou, Fei Zou, and Yuchao Jiang. Scdc: bulk gene expression deconvolution by multiple single-cell rna sequencing references. Briefings in bioinformatics, 22(1):416–427, 2021.

31. Siyao Liu, Aatish Thennavan, Joseph P Garay, JS Marron, and Charles M Perou. Multik: an automated tool to determine optimal cluster numbers in single-cell rna sequencing data. Genome biology, 22(1):1–21, 2021.

32. Luyi Tian, Xueyi Dong, Saskia Freytag, Kim-Anh Lê Cao, Shian Su, Abolfazl JalalAbadi, Daniela Amann-Zalcenstein, Tom S Weber, Azadeh Seidi, Jafar S Jabbari, et al. Benchmarking single cell rna-sequencing analysis pipelines using mixture control experiments. Nature methods, 16(6):479–487, 2019.

33. Ning Leng, Li-Fang Chu, Chris Barry, Yuan Li, Jeea Choi, Xiaomao Li, Peng Jiang, Ron M Stewart, James A Thomson, and Christina Kendziorski. Oscope identifies oscillatory genes in unsynchronized single-cell rna-seq experiments. Nature methods, 12(10):947, 2015.

34. Yohei Sasagawa, Itoshi Nikaido, Tetsutaro Hayashi, Hiroki Danno, Kenichiro D Uno, Takeshi Imai, and Hiroki R Ueda. Quartz-seq: a highly reproducible and sensitive single-cell rna sequencing method, reveals non-genetic gene-expression heterogeneity. Genome biology, 14(4):1–17, 2013.

35. Florian Buettner, Kedar N Natarajan, F Paolo Casale, Valentina Proserpio, Antonio Scialdone, Fabian J Theis, Sarah A Teichmann, John C Marioni, and Oliver Stegle. Computational analysis of cell-to-cell heterogeneity in single-cell rna-sequencing data reveals hidden subpopulations of cells. Nature biotechnology, 33(2):155–160, 2015.

36. Andrew McDavid, Lucas Dennis, Patrick Danaher, Greg Finak, Michael Krouse, Alice Wang, Philippa Webster, Joseph Beechem, and Raphael Gottardo. Modeling bi-modality improves characterization of cell cycle on gene expression in single cells. PLoS computational biology, 10(7):e1003696, 2014.

37. Eduardo Torre, Hannah Dueck, Sydney Shaffer, Janko Gospocic, Rohit Gupte, Roberto Bonasio, Junhyong Kim, John Murray, and Arjun Raj. Rare cell detection by single-cell rna sequencing as guided by single-molecule rna fish. Cell systems, 6(2):171–179, 2018.

38. Sydney M Shaffer, Margaret C Dunagin, Stefan R Torborg, Eduardo A Torre, Benjamin Emert, Clemens Krepler, Marilda Beqiri, Katrin Sproesser, Patricia A Brafford, Min Xiao, et al. Rare cell variability and drug-induced reprogramming as a mode of cancer drug resistance. Nature, 546(7658):431–435, 2017.

39. Solomon Kullback and Richard A Leibler. On information and sufficiency. The annals of mathematical statistics, 22(1):79–86, 1951.

